# Somatostatin Receptor 2 Overexpression in Hepatocellular Carcinoma: Implications for Cancer Biology and Theranostic Applications

**DOI:** 10.1101/2025.01.29.635283

**Authors:** Majid Momeny, Servando Hernandez Vargas, Solmaz AghaAmiri, Jack T. Adams, Tyler M. Bateman, Sukhen C. Ghosh, Belkacem Acidi, Vahid Khalaj, Ahmed O. Kaseb, Hop S. Tran Cao, Ali Azhdarinia

## Abstract

**Background:** Somatostatin receptor 2 (SSTR2) is overexpressed in various tumors, including hepatocellular carcinoma (HCC), yet its role in tumorigenesis remains unclear. This study examines the roles of SSTR2 in the molecular pathology of HCC and explores its potential as a target for SSTR2-directed radiopharmaceuticals in this malignancy.

**Methods:** SSTR2 expression was analyzed across 22 malignancies using TNMplot and specifically in HCC through The Human Protein Atlas. Transcriptomic data, protein expression, and copy number alterations in HCC patients with varying SSTR2 levels were compared using The Cancer Genome Atlas (TCGA). Gene Ontology (GO) enrichment analysis was performed using SRplot, while survival analysis was conducted with GEO datasets.

**Results:** Most HCC patients exhibit moderate levels of SSTR2 expression. Elevated SSTR2 expression is associated with worse overall and disease-specific survival, as well as the activation of pathways involved in tumor growth and metastasis. Furthermore, SSTR2 expression is linked to key oncogenes and receptor tyrosine kinases.

**Conclusions:** SSTR2 in HCC signifies an oncogenic network and represents a promising therapeutic target to inhibit tumor invasion and serve as a theranostic biomarker. HCC patients with elevated SSTR2 expression could benefit from SSTR2-targeted theranostics, enabling enhanced tumor detection and more effective therapy.

## 1. Introduction

Liver cancer, the fourth leading cause of cancer deaths and sixth in new cases globally, is expected to surpass one million cases by 2025 [1]. Hepatocellular carcinoma (HCC) is the most common primary liver malignancy, accounting for approximately 90% of liver cancer cases worldwide [2]. In the United States, HCC incidence has been steadily rising, with an estimated 42,600 new cases and 30,000 deaths in 2023, making it one of the most lethal cancers with a five-year survival rate of around 20% [3]. Major risk factors include chronic hepatitis B and C infections, alcohol-related liver disease, and non-alcoholic fatty liver disease associated with metabolic syndrome. Treatment options for HCC vary based on disease stage and include surgical resection, liver transplantation, locoregional therapies (radiofrequency ablation, transarterial chemoembolization), and systemic therapies, including multi-kinase inhibitors (sorafenib, lenvatinib) and immune checkpoint inhibitors [4]. However, HCC therapy faces significant challenges, including therapy resistance driven by tumor heterogeneity, hypoxia, and alterations in signaling pathways such as PI3K/AKT/mTOR and Wnt/β-catenin [5, 6]. Additionally, a major clinical problem is the lack of reliable biomarkers for early detection and therapeutic response, coupled with our poor understanding of its molecular pathogenesis, which hinders the development of precision therapies [7].

Somatostatin receptor 2 (SSTR2) is one of the five known somatostatin receptor subtypes and is predominantly expressed in various tissues, including the central nervous system, endocrine glands, and gastrointestinal tract. SSTR2 plays a pivotal role in mediating the inhibitory effects of somatostatin on hormone secretion, cell proliferation, and neurotransmission. At the cellular level, SSTR2 is a G protein-coupled receptor (GPCR) that primarily couples with Gi proteins to inhibit adenylyl cyclase activity, leading to decreased cyclic AMP (cAMP) levels. This signaling cascade modulates ion channel activity and inhibits calcium influx, thereby reducing exocytosis of hormones and neurotransmitters. Molecularly, SSTR2 activation also influences downstream pathways such as the MAPK and PI3K/AKT signaling pathways, contributing to its role in regulation of cell cycle progression and apoptosis. Physiologically, SSTR2 regulates critical processes such as insulin and glucagon secretion in pancreatic islets, gastric acid secretion, and intestinal motility [8, 9].

SSTR2 is overexpressed in several human cancers, including neuroendocrine tumors (NETs), small-cell lung carcinomas (SCLCs), and certain gliomas [10–12]. In HCC, nearly 40% of patients exhibit positive SSTR2 membrane staining, with intensities classified as strong (9.6%), moderate (21.2%), and weak (7.7%) [13]. Elevated expression of SSTR2 has been associated with favorable clinical outcomes in various human cancers. In rectal NETs, approximately two-thirds of patients exhibit SSTR2 expression, which correlates with smaller tumor size, lower tumor stage, and improved overall survival [14]. Similar correlation has been found in gliomas, where high SSTR2 levels are linked to lower WHO grades and better prognoses [12]. Conversely, high SSTR2 expression has been associated with poor prognosis in nasopharyngeal carcinoma [15]. Similarly, in SCLC, SSTR2 signaling has been linked to tumor progression [11], suggesting that SSTR2 may exhibit both oncogenic and tumor suppressor functions depending on the cancer type.

Overexpression of SSTR2 in NETs has permitted to development of clinically approved theranostics, a strategy that combines non-invasive nuclear imaging with targeted radionuclide delivery to improve tumor detection and treatment efficacy [16]. Radiolabeled somatostatin analogs such as ^68^Ga-DOTATATE enable positron emission tomography (PET) imaging for tumor detection, disease staging, and treatment monitoring, while peptide receptor radioligand therapy (PRRT) with ^177^Lu-DOTATATE enables precision treatment of tumors with cell-damaging beta particles, which minimizes damage to healthy tissues and reduces disease progression or mortality by 80% [17]. The potential use of a similar strategy for managing other SSTR2-expressing cancers, including HCC, remains largely unexplored.

Although SSTR2 is significantly expressed in HCC, its relationship with the clinicopathological features of this cancer remains poorly characterized. Furthermore, the functional roles of SSTR2 in the molecular pathology of HCC and the signaling pathways it regulates are not yet fully understood. This study aims to provide preliminary evidence for the clinical relevance of SSTR2 in HCC and to investigate its potential as both a therapeutic and theranostic target in this malignancy.

## 2. Materials and Methods

The publicly available dataset TNMplot was utilized to analyze the expression of *SSTR2* in normal versus tumor tissues across 22 human malignancies. The expression of SSTR2 in HCC patients was investigated using The Human Protein Atlas Dataset (https://www.proteinatlas.org/). The Cancer Genome Atlas (TCGA) dataset, which provides comprehensive genomic data and molecular features for approximately 33 cancer types [18], was used to compare transcriptomic profiling, protein expression, and arm-level copy number alterations (CNAs) between HCC patients with high and low *SSTR2* expression. RNA-sequencing data from the TCGA were graphed using the GO enrichment analysis from the SRplot database [19]. Patient survival data were obtained from pooled Gene Expression Omnibus datasets via the Kaplan–Meier plot [20]. Patients were categorized into two groups based on the upper and lower quartiles of normalized *SSTR2* expression. The Kaplan–Meier method was used to compute overall survival, relapse-free survival, progression-free survival, and disease-specific survival.

## 3. Results

Using the web tool TNMplot, we assessed *SSTR2* mRNA levels across 22 different cancer types. Our results reveal that *SSTR2* levels are significantly higher in tumor tissues compared to normal tissues in several malignancies, including liver cancer (Supplementary Fig. 1). Furthermore, data from The Human Protein Atlas Database indicates that most HCC patients express SSTR2 protein. Immunohistochemistry (IHC) staining for SSTR2 in 12 HCC patients (using antibody #HPA007264 from Sigma) showed medium staining in 9 patients, weak staining in 2 patients, and no staining in 1 patient (Fig. 1A). These findings are consistent with previous studies, which reported membrane staining of SSTR2 in about 40% of HCC cases, with 9.6% showing strong staining, 21.2% moderate staining, and 7.7% weak staining [13].

**Figure 1.**
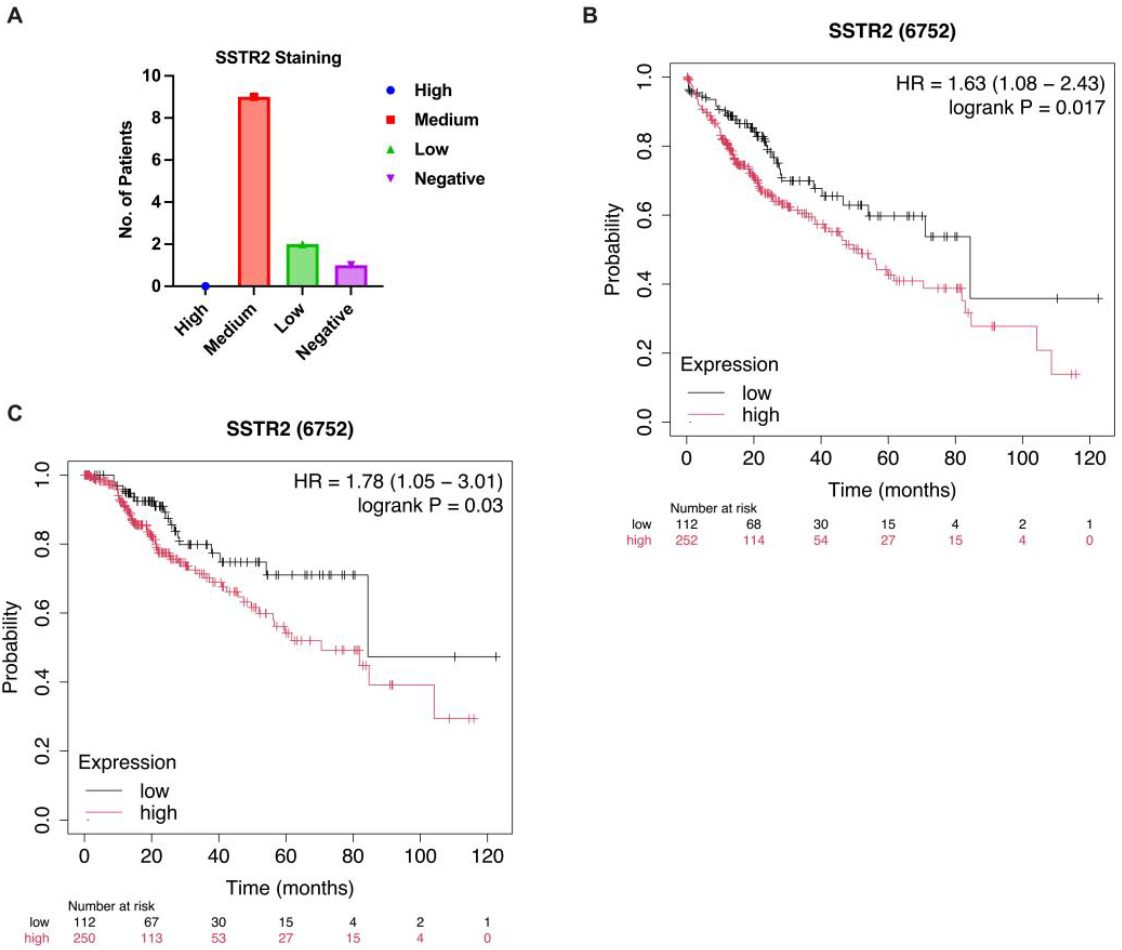
(A) Immunohistochemical (IHC) staining for SSTR2 in 12 hepatocellular carcinoma (HCC) patients demonstrates moderate staining in most cases. Data were sourced from the Human Protein Atlas (https://www.proteinatlas.org). (B, C) Elevated SSTR2 expression in HCC is associated with poorer overall survival and disease-specific survival, as shown by Kaplan-Meier survival analysis.

Due to the higher expression of *SSTR2* in liver cancer tissues compared to normal tissues (Supplementary Fig. 1), we hypothesized that it may play roles in HCC tumorigenesis. Consistent with this, using the Kaplan Meier plotter, we discovered that higher expression of SSTR2 in HCC patients predicts both poor overall and disease-specific survival (Fig. 1B, C). Furthermore, dividing the patients from the TCGA HCC dataset [18] based on *SSTR2* mRNA expression levels measured by RNA sequencing into SSTR2^high^ (LogFC>1, FDR<0.05, n=60) and SSTR2^low^ (LogFC<−1, FDR<0.05, n=59) groups shows that higher *SSTR2* expression predicts poorer overall survival (Log-rank Test *p* value = 0.0393) (Supplementary Fig. 2). Moreover, analysis of the RNA-sequencing data from these two groups shows that the expression of ~1800 genes is significantly (logFC>1, *p* value <0.05) higher in SSTR2^high^ patients compared to SSTR2^low^ group, whereas ~500 gene exhibit a lower expression (logFC<−1, *p* value <0.05) (Fig. 2A). Gene Set Enrichment Analysis (GSEA) analysis of the genes with a higher expression in the SSTR2^high^ group revealed activation of hallmark pathways including chemokine signaling, PI3K/AKT pathway, cell adhesion and tumor-stroma interaction (Fig. 2B).

**Figure 2.**
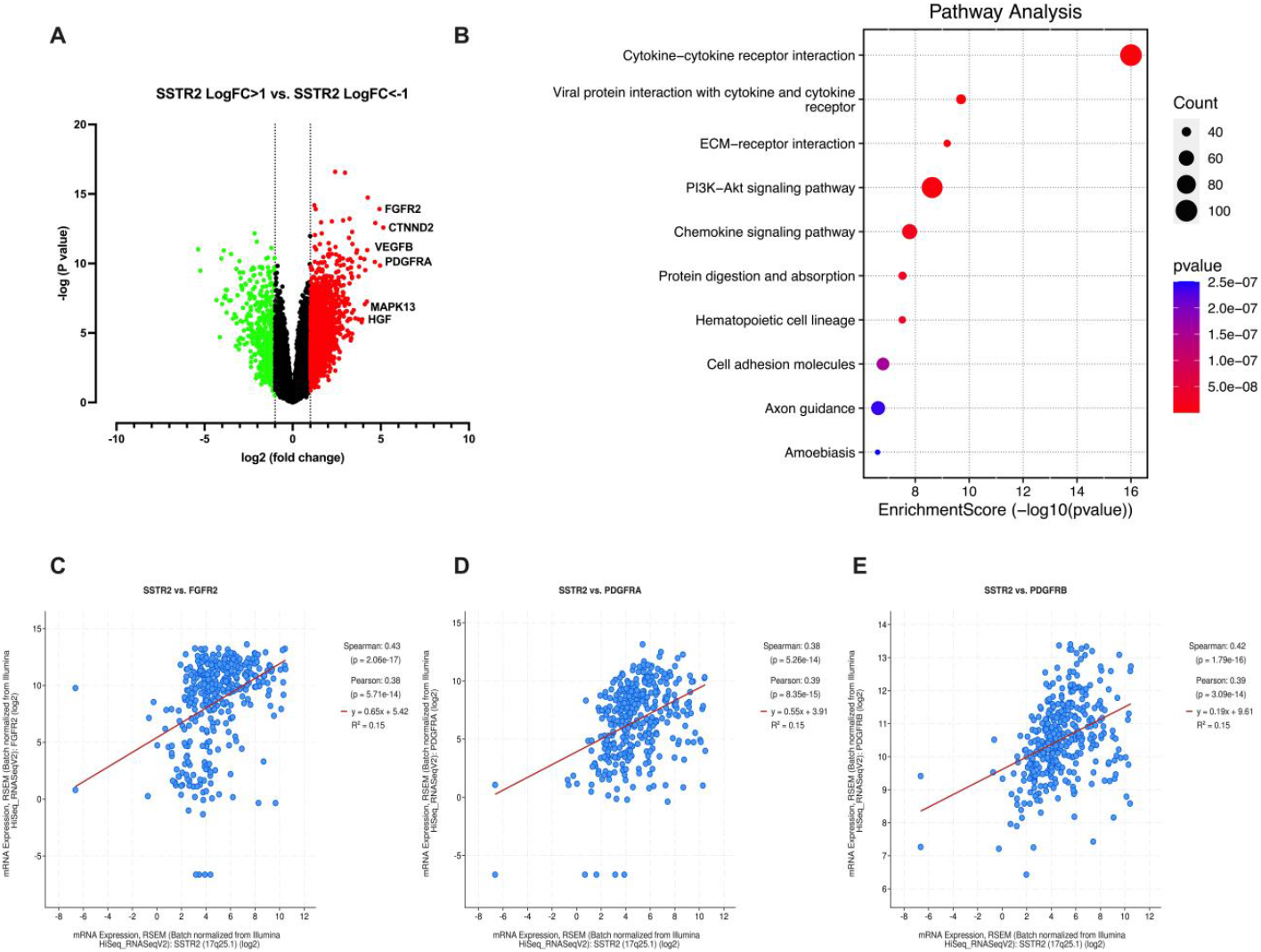
(A) Patients from the TCGA HCC dataset [18] were stratified into SSTR2^high^ (LogFC > 1, FDR < 0.05, n = 60) and SSTR2^low^ (LogFC < −1, FDR < 0.05, n = 59) groups based on *SSTR2* mRNA expression levels. RNA-sequencing analysis revealed significant differences in the expression of key HCC oncogenes between the two groups. (B) Gene Set Enrichment Analysis (GSEA) of genes upregulated in the SSTR2^high^ group highlighted the activation of hallmark pathways, including chemokine signaling, the PI3K/AKT pathway, cell adhesion, and tumor-stroma interactions. (C-E) In TCGA HCC patients [18], *SSTR2* mRNA levels positively correlated with key receptor tyrosine kinases, such as *FGFR2, PDGFRA*, and *PDGFRB*.

Delta-catenin (CTNND2), vascular endothelial growth factor B (VEGFB), platelet-derived growth factor receptor alpha (PDGFRA), mitogen-activated protein kinase 13 (MAPK13), and hepatocyte growth factor (HGF) have higher expression in the SSTR2^high^ group (Fig. 2A). We also observed a positive correlation between the mRNA levels of *SSTR2* and key receptor tyrosine kinases, including *PDGFRA*, platelet-derived growth factor receptor beta (*PDGFRB*) and fibroblast growth factor receptor 2 (*FGFR2*) in the TCGA HCC patients [18] (Fig. 2C-E). VEGFB, PDGFRα, MAPK13, and HGF promote tumor growth, survival, angiogenesis, and metastasis, and their elevated expression is linked to poor prognosis in HCC [21]. Delta-catenin, part of the p120-catenin family, is key in cell adhesion and cancer metastasis, especially promoting epithelial-mesenchymal transition (EMT) [22], a process that facilitates cancer cell migration and invasion [23].

We compared protein expression between the two patient groups and found that several oncogenic proteins, including plasminogen activator inhibitor-1 (PAI-1), TP53-induced glycolysis and apoptosis regulator (TIGAR), spleen tyrosine kinase (SYK), fibronectin 1, cyclin B1, MAPK1, fatty acid synthase (FASN), and SRC, were significantly elevated in the SSTR2^high^ group (Fig. 3A). PAI-1, involved in matrix dissolution and cancer metastasis, is linked to increased invasion and poor prognosis in HCC [24]. TIGAR regulates cell growth, and its knockdown induces apoptosis and autophagy in HCC cells [25]. SYK overexpression correlates with EMT, metastasis and vascular invasion in HCC [26]. Fibronectin induces EMT in HCC, while cyclin B1, MAPK1, and SRC are associated with aggressive tumor behavior and poor prognosis by affecting tumor growth and metastasis [27–30].

**Figure 3.**
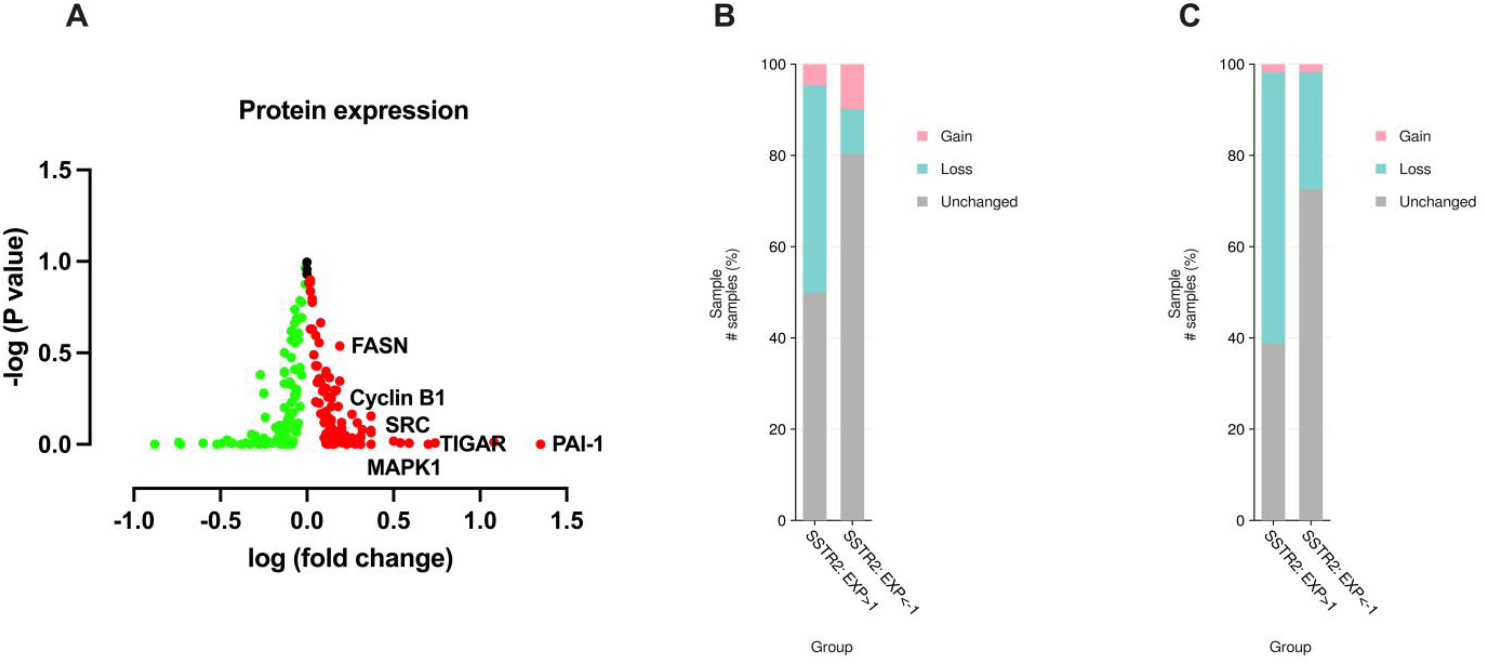
(A) Protein expression analysis between SSTR2^high^ and SSTR2^low^ TCGA HCC patients [18] revealed significantly elevated levels of oncogenic proteins, including PAI-1, TIGAR, SYK tyrosine kinase, fibronectin 1, cyclin B1, MAPK1, fatty acid synthase, and SRC, in the SSTR2^high^ group. These proteins are pivotal in promoting cell survival, epithelial-to-mesenchymal transition (EMT), and tumor invasion in HCC. (B, C) Arm-level copy number alteration analysis showed a higher frequency of 1p and 16q losses in SSTR2^high^ compared to SSTR2^low^ patients. Key tumor suppressors, including *CDH1* (1p22.1) and *RUNX3* (16q24.1), are implicated in inhibiting tumor invasion and metastasis by suppressing EMT.

We then analyzed arm-level copy number alterations (CNAs) between the two patient groups and found significantly higher rates of 1p and 16q losses in SSTR2^high^ patients compared to SSTR2^low^ group (Fig. 3B, C). *CDH1* at 1p22.1 encodes E-cadherin, crucial for cell adhesion and metastasis suppression [31]. *RUNX3* at 16q24.1 acts as a tumor suppressor, promoting E-cadherin expression and inhibiting EMT and cell invasion in HCC [32]. Our findings suggest that higher *SSTR2* expression in linked with a more invasive and aggressive phenotype in HCC patients, providing a molecular explanation for the contribution of SSTR2 to the poor clinical outcomes in this malignancy.

## 4. Discussion

Our study suggests that high SSTR2 expression in HCC could significantly impact its clinical management, serving as both a prognostic biomarker for patient outcomes and a theranostic biomarker for diagnostic and treatment purposes. Our novel observations suggest a potential link between SSTR2 expression, receptor tyrosine kinase expression, tumor metastasis, clinical outcomes, and therapeutic sensitivity.

The VEGFR, PDGFR, and RAF/MEK/ERK signal transduction pathways are integral to the pathogenesis and progression of HCC. VEGFRs are key regulators of angiogenesis, which facilitates tumor growth and metastasis by promoting blood vessel formation in HCC [33]. PDGFRs play crucial roles in tumor-stroma interactions, contributing to HCC progression through enhanced stromal support, angiogenesis, and fibrogenesis [34, 35]. Additionally, the RAF/MEK/ERK pathway, a central mediator of the MAPK signaling cascade, is often dysregulated in HCC, driving cellular proliferation, survival, and resistance to apoptosis [30]. Dysregulation of these pathways promotes HCC tumorigenesis and their blockade, either individually or in combination, holds promise as a therapeutic strategy to inhibit tumor growth and malignant progression in HCC [36].

Sorafenib, an FDA-approved multi-kinase inhibitor for HCC, exerts its anti-tumor effects by inhibiting cell proliferation and angiogenesis through targeting the VEGFR, PDGFR, and RAF/MEK/ERK pathways [37]. The findings of this study reveal a positive correlation between high SSTR2 expression and the upregulation of these key oncogenic pathways in HCC. However, whether SSTR2 directly regulates these pathways or whether they collectively contribute to an oncogenic network in HCC remains an open and intriguing question. Addressing this question could unveil new opportunities for prognosis and therapeutic intervention. SSTR2 may play a role in the molecular pathology of HCC by regulating these oncogenic pathways, making it a promising novel therapeutic target for the treatment of HCC. Moreover, targeting SSTR2 with specific modulators may disrupt the VEGFR, PDGFR, and RAF/MEK/ERK pathways, offering a synergistic approach with sorafenib to inhibit tumor growth and angiogenesis more effectively.

Despite its broad spectrum of molecular targets, the clinical efficacy of sorafenib in HCC remains limited. Clinical trials report that only approximately 30% of patients derive substantial benefit, with a median overall survival improvement of just 2–3 months. Moreover, resistance to sorafenib frequently develops within six months of treatment initiation, underscoring the roles of both intrinsic and acquired resistance mechanisms [38, 39]. These limitations have spurred significant research efforts to enhance the therapeutic effectiveness of sorafenib. Combination therapies have emerged as promising strategies to overcome resistance and achieve synergistic anti-tumor effects [24]. While the role of SSTR2 in mediating resistance to sorafenib in HCC remains to be elucidated, SSTR2-targeted theranostic applications have significant potential to synergize with sorafenib and improve its efficacy in sorafenib-resistant SSTR2-positive cases. Supporting this hypothesis, our recent findings demonstrate that ^177^Lu-DOTATATE enhances the anti-tumor activity of sorafenib in an HCC cell line [40].

Another noteworthy finding of this study is the elevated expression of key proteins involved in extracellular matrix degradation, motility, and invasion in HCC patients with higher *SSTR2* expression, including PAI-1, TIGAR, SYK, fibronectin, MAPK1, FASN, and SRC. Consistently, we observed a higher frequency of deletions in *CDH1*, which encodes E-cadherin, a critical suppressor of tumor metastasis [31], and *RUNX3*, which promotes E-cadherin expression and inhibits EMT [32], in these patients. The loss of E-cadherin and activation of pro-metastatic pathways, including SRC and MAPK, further accelerates invasion and metastasis in HCC [41–43]. Metastasis, a critical driver of poor prognosis in HCC, involves complex processes such as EMT, ECM degradation, angiogenesis, and immune evasion. These mechanisms facilitate tumor dissemination to distant organs, such as the lungs and lymph nodes, and correlate with aggressive tumor behavior, therapy resistance, and reduced survival [44]. Collectively, our findings suggest that SSTR2 may promote cell motility and invasion in HCC by regulating pro-metastatic pathways and EMT. These insights highlight the importance of further investigations to elucidate the role of SSTR2 in cancer metastasis in HCC and to explore its potential as a therapeutic target to inhibit tumor invasion in this highly fatal malignancy. These preliminary results warrant further validation through advanced animal models and early-phase clinical trials to explore their translational potential.

## 5. Conclusions

Our findings indicate that SSTR2 may be part of an oncogenic network that promotes malignant progression in HCC, warranting further investigation into its roles in this malignancy. HCC patients with elevated SSTR2 expression are likely to benefit from ^68^Ga-DOTATATE PET/CT, which could improve tumor detection, assess therapeutic responses to systemic and immunotherapies, guide patient selection for PRRT therapy, and monitor disease progression. Additionally, SSTR2-positive HCC patients may benefit from therapy with ^177^Lu-DOTATATE. Future research should prioritize the efficacy of SSTR2-based imaging and therapies for liver metastases with high SSTR2 expression, potentially leading to more precise treatments and improved patient outcomes. Additionally, insights from this study may inform novel treatment combinations, particularly with emerging SSTR2-targeted alpha therapies, which have shown exceptional efficacy in NETs [45] and could be applicable to HCC treatment. Further research is essential to validate these hypotheses and clarify the clinical utility of SSTR2 as a therapeutic target and a theranostic biomarker in HCC.

## Supplementary Materials

The following supporting information can be downloaded at: www.mdpi.com/xxx/s1, Figure S1: Higher expression of *SSTR2* in human malignancies compared to normal tissues; Figure S2: Higher expression of *SSTR2* predicts a poor overall survival in HCC patients.

## Author Contributions

All authors contributed to the study conception and design. The first draft of the manuscript was written by Majid Momeny and all authors commented on previous versions of the manuscript. The study was supervised by Majid Momeny and Ali Azhdarinia. All authors read and approved the final manuscript.

## Institutional Review Board Statement

Not applicable.

## Informed Consent Statement

Not applicable.

## Data Availability Statement

Data from this study are available upon reasonable request from the corresponding authors.

## Abbreviations

The following abbreviations are used in this manuscript:

^68^Ga: Gallium-68
HCC: Hepatocellular carcinoma
^177^Lu: Lutetium-177
SSTR2: Somatostatin receptor 2

## References

1. Brown, Z.J., et al., Management of Hepatocellular Carcinoma: A Review. JAMA Surg, 2023. 158(4): p. 410–420.

2. Craig, A.J., et al., Tumour evolution in hepatocellular carcinoma. Nat Rev Gastroenterol Hepatol, 2020. 17(3): p. 139–152.

3. Siegel, R.L., et al., Cancer statistics, 2023. CA Cancer J Clin, 2023. 73(1): p. 17–48.

4. Finn, R.S., et al., Atezolizumab plus Bevacizumab in Unresectable Hepatocellular Carcinoma. N Engl J Med, 2020. 382(20): p. 1894–1905.

5. Safri, F., et al., Heterogeneity of hepatocellular carcinoma: from mechanisms to clinical implications. Cancer Gene Ther, 2024. 31(8): p. 1105–1112.

6. Marin, J.J.G., et al., Molecular Bases of Drug Resistance in Hepatocellular Carcinoma. Cancers (Basel), 2020. 12(6).

7. Llovet, J.M., et al., Immunotherapies for hepatocellular carcinoma. Nat Rev Clin Oncol, 2022. 19(3): p. 151–172.

8. Reubi, J.C., et al., Somatostatin receptor sst1-sst5 expression in normal and neoplastic human tissues using receptor autoradiography with subtype-selective ligands. Eur J Nucl Med, 2001. 28(7): p. 836–46.

9. Gunther, T., et al., International Union of Basic and Clinical Pharmacology. CV. Somatostatin Receptors: Structure, Function, Ligands, and New Nomenclature. Pharmacol Rev, 2018. 70(4): p. 763–835.

10. Guenter, R., et al., Overexpression of somatostatin receptor type 2 in neuroendocrine tumors for improved Ga68-DOTATATE imaging and treatment. Surgery, 2020. 167(1): p. 189–196.

11. Lehman, J.M., et al., Somatostatin receptor 2 signaling promotes growth and tumor survival in small-cell lung cancer. Int J Cancer, 2019. 144(5): p. 1104–1114.

12. He, J.H., et al., SSTR2 is a prognostic factor and a promising therapeutic target in glioma. Am J Transl Res, 2021. 13(10): p. 11223–11234.

13. Lequoy, M., et al., Somatostatin receptors in resected hepatocellular carcinoma: status and correlation with markers of poor prognosis. Histopathology, 2017. 70(3): p. 492–498.

14. Kim, J.Y., et al., Somatostatin receptor 2 (SSTR2) expression is associated with better clinical outcome and prognosis in rectal neuroendocrine tumors. Sci Rep, 2024. 14(1): p. 4047.

15. Xu, Y., et al., SSTR2 positively associates with EGFR and predicts poor prognosis in nasopharyngeal carcinoma. J Clin Pathol, 2024. 77(12): p. 829–834.

16. Palekar-Shanbhag, P., et al., Theranostics for cancer therapy. Curr Drug Deliv, 2013. 10(3): p. 357–62.

17. Strosberg, J., et al., Phase 3 Trial of (177)Lu-Dotatate for Midgut Neuroendocrine Tumors. N Engl J Med, 2017. 376(2): p. 125–135.

18. Cerami, E., et al., The cBio cancer genomics portal: an open platform for exploring multidimensional cancer genomics data. Cancer Discov, 2012. 2(5): p. 401–4.

19. Tang, D., et al., SRplot: A free online platform for data visualization and graphing. PLoS One, 2023. 18(11): p. e0294236.

20. Lanczky, A. and B. Gyorffy, Web-Based Survival Analysis Tool Tailored for Medical Research (KMplot): Development and Implementation. J Med Internet Res, 2021. 23(7): p. e27633.

21. Dimri, M. and A. Satyanarayana, Molecular Signaling Pathways and Therapeutic Targets in Hepatocellular Carcinoma. Cancers (Basel), 2020. 12(2).

22. Zhang, H., et al., Delta-catenin promotes the proliferation and invasion of colorectal cancer cells by binding to E-cadherin in a competitive manner with p120 catenin. Target Oncol, 2014. 9(1): p. 53–61.

23. Brabletz, T., et al., EMT in cancer. Nat Rev Cancer, 2018. 18(2): p. 128–134.

24. Zheng, Q., et al., Invasion and metastasis of hepatocellular carcinoma in relation to urokinase-type plasminogen activator, its receptor and inhibitor. J Cancer Res Clin Oncol, 2000. 126(11): p. 641–6.

25. Ye, L., et al., Knockdown of TIGAR by RNA interference induces apoptosis and autophagy in HepG2 hepatocellular carcinoma cells. Biochem Biophys Res Commun, 2013. 437(2): p. 300–6.

26. Hong, J., et al., Expression of variant isoforms of the tyrosine kinase SYK determines the prognosis of hepatocellular carcinoma. Cancer Res, 2014. 74(6): p. 1845–56.

27. Zhang, L., et al., Fibronectin 1 derived from tumor-associated macrophages and fibroblasts promotes metastasis through the JUN pathway in hepatocellular carcinoma. Int Immunopharmacol, 2022. 113(Pt A): p. 109420.

28. Lv, S., et al., Inhibition of cyclinB1 Suppressed the Proliferation, Invasion, and Epithelial Mesenchymal Transition of Hepatocellular Carcinoma Cells and Enhanced the Sensitivity to TRAIL-Induced Apoptosis. Onco Targets Ther, 2020. 13: p. 1119–1128.

29. Zhao, P.W., et al., SRC-1 and Twist1 are prognostic indicators of liver cancer and are associated with cell viability, invasion, migration and epithelial-mesenchymal transformation of hepatocellular carcinoma cells. Transl Cancer Res, 2020. 9(2): p. 603–612.

30. Moon, H. and S.W. Ro, MAPK/ERK Signaling Pathway in Hepatocellular Carcinoma. Cancers (Basel), 2021. 13(12).

31. Na, T.Y., et al., The functional activity of E-cadherin controls tumor cell metastasis at multiple steps. Proc Natl Acad Sci U S A, 2020. 117(11): p. 5931–5937.

32. Tanaka, S., et al., Runt-related transcription factor 3 reverses epithelial-mesenchymal transition in hepatocellular carcinoma. Int J Cancer, 2012. 131(11): p. 2537–46.

33. Morse, M.A., et al., The Role of Angiogenesis in Hepatocellular Carcinoma. Clin Cancer Res, 2019. 25(3): p. 912–920.

34. Stock, P., et al., Platelet-derived growth factor receptor-alpha: a novel therapeutic target in human hepatocellular cancer. Mol Cancer Ther, 2007. 6(7): p. 1932–41.

35. Zhang, T., et al., Overexpression of platelet-derived growth factor receptor alpha in endothelial cells of hepatocellular carcinoma associated with high metastatic potential. Clin Cancer Res, 2005. 11(24 Pt 1): p. 8557–63.

36. Llovet, J.M., et al., Molecular pathogenesis and systemic therapies for hepatocellular carcinoma. Nat Cancer, 2022. 3(4): p. 386–401.

37. Wilhelm, S.M., et al., Preclinical overview of sorafenib, a multikinase inhibitor that targets both Raf and VEGF and PDGF receptor tyrosine kinase signaling. Mol Cancer Ther, 2008. 7(10): p. 3129–40.

38. Cheng, A.L., et al., Efficacy and safety of sorafenib in patients in the Asia-Pacific region with advanced hepatocellular carcinoma: a phase III randomised, double-blind, placebo-controlled trial. Lancet Oncol, 2009. 10(1): p. 25–34.

39. Llovet, J.M., et al., Sorafenib in advanced hepatocellular carcinoma. N Engl J Med, 2008. 359(4): p. 378–90.

40. Momeny, M., et al., SSTR2-targeted theranostics in hepatocellular carcinoma. Cancers, 2025. 17(2): p. 162.

41. Hashiguchi, M., et al., Clinical implication of ZEB-1 and E-cadherin expression in hepatocellular carcinoma (HCC). BMC Cancer, 2013. 13: p. 572.

42. Zhao, S., et al., The role of c-Src in the invasion and metastasis of hepatocellular carcinoma cells induced by association of cell surface GRP78 with activated alpha2M. BMC Cancer, 2015. 15: p. 389.

43. Mo, S., et al., Down regulated oncogene KIF2C inhibits growth, invasion, and metastasis of hepatocellular carcinoma through the Ras/MAPK signaling pathway and epithelial-to-mesenchymal transition. Ann Transl Med, 2022. 10(3): p. 151.

44. Llovet, J.M., et al., Hepatocellular carcinoma. Nat Rev Dis Primers, 2021. 7(1): p. 6.

45. Han, G., et al., RYZ101 (Ac-225 DOTATATE) Opportunity beyond Gastroenteropancreatic Neuroendocrine Tumors: Preclinical Efficacy in Small-Cell Lung Cancer. Mol Cancer Ther, 2023. 22(12): p. 1434–1443.

