## Supplementary figures and images for "Somatostatin Receptor 2 Overexpression in Hepatocellular Carcinoma: Implications for Cancer Biology and Theranostic Applications"

### Supplementary Fig 1 and 2

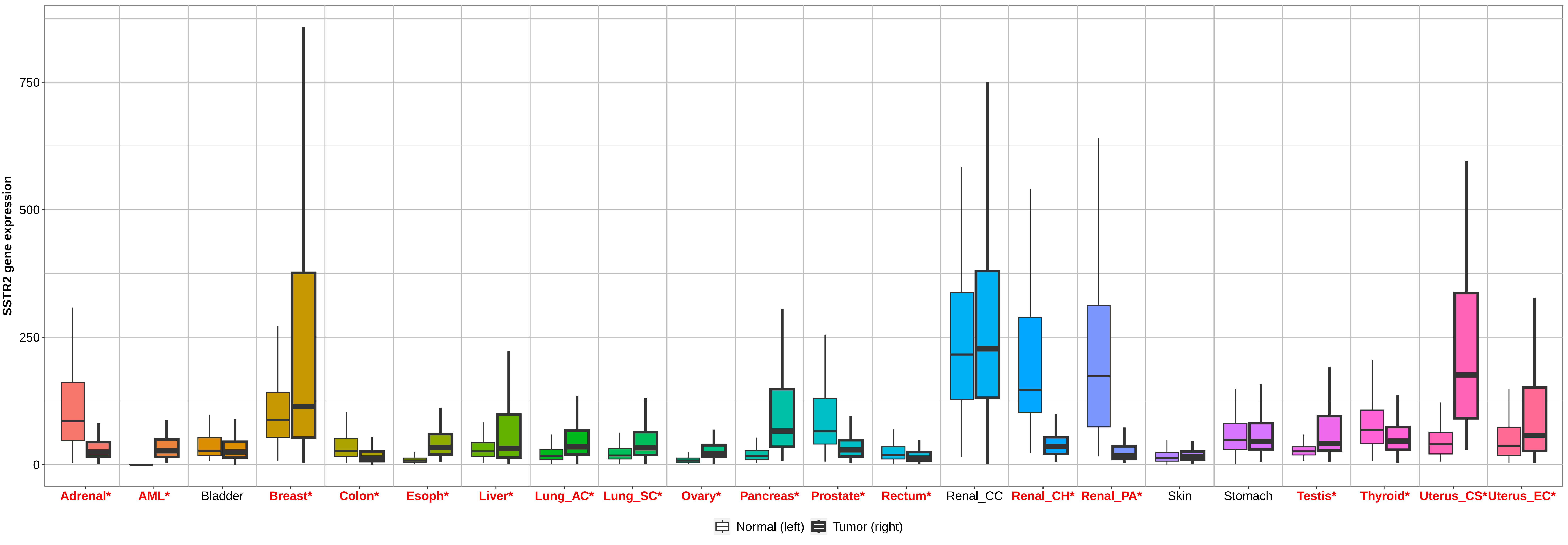

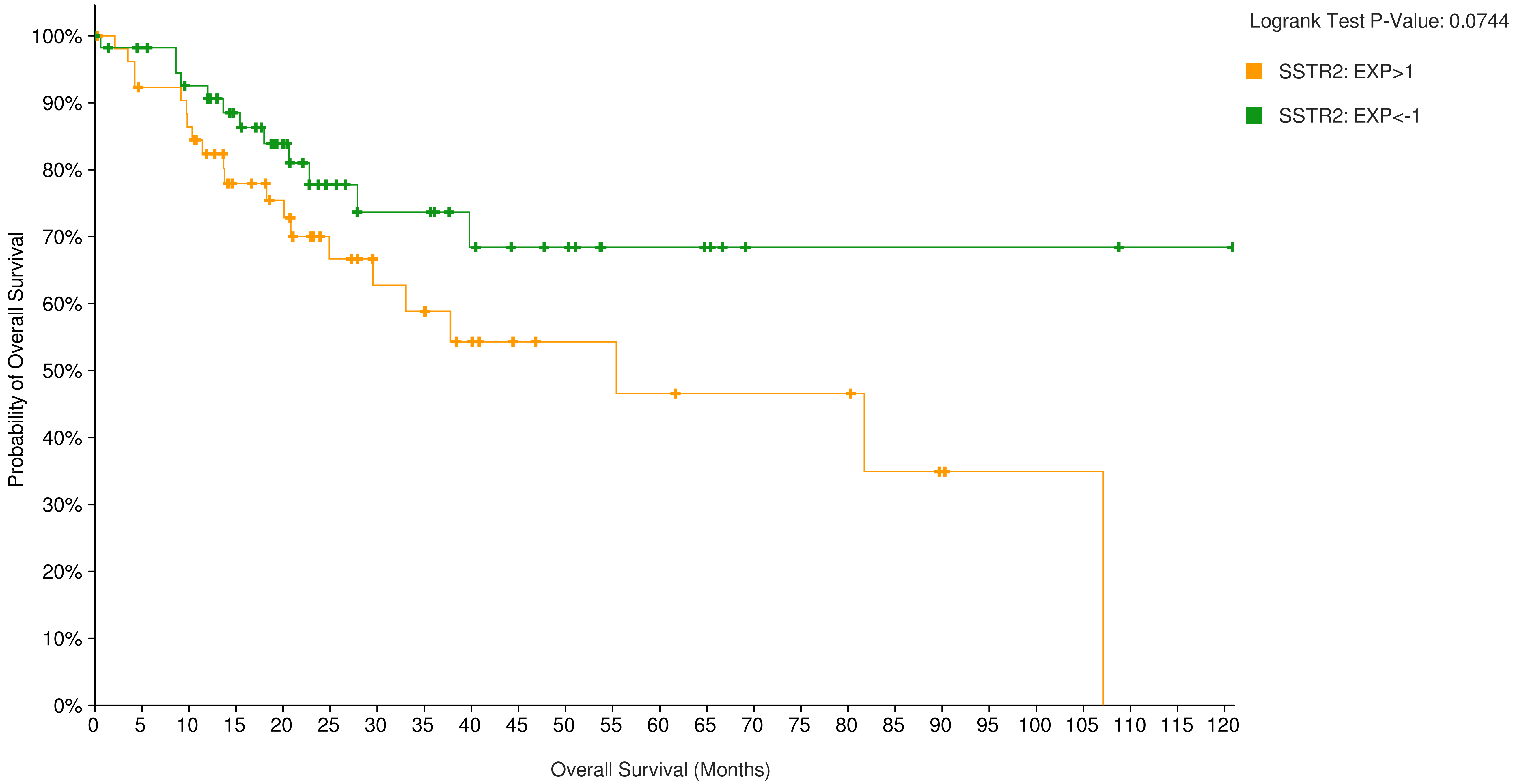


**Number at risk (n)**

| | | | | | | | | | | | | | | | | | | | | | | | | | |
|---|---|---|---|---|---|---|---|---|---|---|---|---|---|---|---|---|---|---|---|---|---|---|---|---|---|
| SSTR2: EXP>1 | 56 | 47 | 44 | 33 | 29 | 20 | 16 | 15 | 11 | 8 | 7 | 7 | 6 | 5 | 5 | 5 | 5 | 3 | 2 | 1 | 1 | 1 | 0 | 0 | 0 |
| SSTR2: EXP<-1 | 57 | 53 | 48 | 40 | 30 | 21 | 17 | 17 | 13 | 11 | 10 | 6 | 6 | 5 | 2 | 2 | 2 | 2 | 2 | 2 | 2 | 2 | 1 | 1 | 1 |
